# Integrated RNA-seq and sRNA-seq revealed differences in transcriptome between susceptible and resistant tomato responding to *Fusarium oxysporum*

**DOI:** 10.1101/324574

**Authors:** Min Zhao, Hui-Min Ji, Ying Gao, Xin-Xin Cao, Hui-Ying Mao, Peng Liu, Shou-Qiang Ouyang

**Affiliations:** College of Horticulture and Plant Protection, Yangzhou University, 48 East Wenhui Road, Yangzhou, Jiangsu, 225009, China; Texting Center, Yangzhou University, 48 East Wenhui Road, Yangzhou, Jiangsu, 225009, China; Joint International Research Laboratory of Agriculture and Agri-Product Safety, Yangzhou University, 48 East Wenhui Road, Yangzhou, Jiangsu, 225009, China

**Keywords:** Tomato, *Fusarium oxysporum* f. sp. *Lycopersici*, Transcriptome, Wilt disease, Resistance

## Abstract

Tomato wilt disease caused by *Fusarium oxysporum* f. sp. *lycopersici* (FOL) is a worldwide destructive disease of tomato. As exploring gene expression and function approaches constitute an initial point for investigating pathogen-host interaction, we performed RNA-seq and sRNA-seq analysis to unravel regulated genes and miRNAs in tomato infected by FOL. Differentially expressed (DE) protein coding gene and miRNA gene profiles upon inoculation with FOL were presented at twenty-four hours post-inoculation including four treatments. Total of more than 182.6 million and 132.2 million high quality clean reads were obtained by RNA-seq and sRNA-seq, respectively. A large overlap was found in DE mRNAs between susceptible cultivar Moneymaker and resistant cultivar Motelle. All Gene Ontology terms were mainly classified into catalytic activity, metabolic process and binding. Combining with qRT-PCR, five disease resistance genes, Solyc01g095630, Solyc03g059080, Solyc00g174340, Solyc11g071750 and Solyc05g050350, were verified to involved in the disease resistance in the resistant cultivar Motelle treated with FOL. Northern blot analysis further confirmed the results from sRNA-Seq and demonstrated that several miRNAs including Sly-miR477-5p, sly-miR167a, novel_mir_675, novel_mir_504 and novel_mir_762 conferred FOL infection. Our data resulted that pathogen resistant genes/miRNAs may play a critical role with the benefit of a coordinated machinery in prompting the response in prompting FOL response in tomato, which offered us with a future direction and surely help in generating models of mediated resistance responses with assessment of genomic gene expression patterns.

---

*Fusarium oxysporum* f. sp. *lycopersici* (race 2) (as FOL in this study) is a biotrophic pathogen which is the causal agent of tomato wilt. *Fusarium oxysporum* is one of the majority destructive necrotrophic pathogens affecting a large species complex including tomato by causing wilt disease worldwide (Di *et al.* 1998; Leslie *et al.* 2006). Under appropriate conditions, FOL infection leads to clogged vessels resulting in yellowing of leaves, wilting and finally death of the whole plant. According to their specific pathogenicity to tomato cultivars, three physiological races of FOL are distinguished (Di *et al.* 1998; Leslie *et al.* 2006; Takken *et al.* 2010; Kawabe *et al.* 2005.)

Tomato (*Solanum lycopersicum*) is a worldwide agriculture economic crop and also has been studied as a crucial model plant for studying the genetics and molecular basis of resistance mechanisms. Four plant resistance (*R*) genes have been discovered from wild tomato species including the *I* and *I-2* genes from *S. pimpinellifolium*, and the *I-3* and *I-7* gene from *S. pennellii.* Among these four *R* genes, so far, *I-2, I-3* and *I-7* have been cloned, encode an NBS-LRR protein like most known *R* genes (Ori *et al.* 1997; Simons *et al.* 1998; Catanzariti *et al.* 2015; Gonzalez-Cendales *et al.* 2016). Previous works have unveiled that the *I-2* and *I-3* gene confer resistance to race 2 and race 3 strains of FOL, respectively (Simons *et al.* 1998; Catanzariti *et al.* 2015). The *I-2* locus encodes an *R* protein that recognizes avr2 from FOL (race 2) (Houterman *et al.* 2009). The *I-3* encodes an S-receptor-like kinase (SRLK) genes that confers *Avr3*-dependent resistance to FOL (race 3) (Catanzariti *et al.* 2015). Previously, two near-isogenic tomato cultivars susceptible Moneymaker (*i-2*/*i-2*) and resistant Motelle (*I-2*/*I-2*) were recruited to study the interaction between tomato and FOL. The genotypes of these two tomato cultivars are for *I-2* and respond to FOL infection (Di *et al.* 1998; De *et al.* 2001; Yu *et al.* 2008). 3210110901201000026083

Basically, transcriptome analysis is a very important tool to discover the molecular basis of plant-pathogen interaction globally, allowing dissection of the pattern of pathogen activities and molecular repertoires available for defense responses in host plant. By taking advantage of RNA-seq technology, certain number of transcriptome profiling studies of plants inoculated by *Fusarium* fungus have been presented including banana (Guo *et al.* 2014), cabbage (Xing *et al.* 2016), watermelon (Liu *et* al.2015), mango (Liu *et al.* 2016), and *Arabidopsis* (Chen *et al.* 2014; Gupta *et al.* 2014). Upon to pathogens infection, plants activate a few of defense responses to resistant diseases caused by according pathogens, such as hypersensitive reaction (HR), structural alterations, reactive oxygen species (ROS) accumulation, synthesis of secondary metabolites and defense molecules (Park *et al.* 2003; Shah *et al.* 2003; Ros *et al.* 2004).

MicroRNAs (miRNAs) (20-24 nucleotides in length) are derived from endogenous single-stranded non-coding small RNAs with imperfectly base-paired hairpin structures, which regulate gene expression at the post-transcriptional level via base-pairing cleavage or translational repression of recognized target mRNAs (Llave *et al.* 2002; Reinhart *et al.* 2000; Llave *et al.* 2002; Brodersen *et al.* 2008). To date, erupting studies have demonstrated that miRNAs play critical roles in various biotic and abiotic stress responses, especially stress responses (Sunkar *et al.* 2007; Sunkar *et al.* 2006) and innate immunity (Padmanabhan *et al.* 2009; Qiao *et al.* 2013; Weiberg *et al.* 2013). MiRNAs might confer pathogen response by regulating plant hormonal network, such as auxin, jasmonic acid (JA), ethylene (ET), as well as salicylic acid (SA)-mediated defense. For example, miR393 acts as a positive regulator by target the auxin-related F box/auxin receptor transport inhibitor response 1 (AFB/TIR1), which results in enhancing accumulation of pathogen-resistant protein (Robert-Seilaniantz *et al.* 2011; Wong *et al.* 2011). Our previous study showed that two miRNAs, sly-miR482f and sly-miR5300, were verified to play a crucial role in response to FOL infection in tomato (Ouyang *et al.* 2014). However, there was still no other reported miRNAs conferring tomato wilt disease so far. In addition, there is far less attention to understand how genes and miRNAs are integrated into the dynamic and complex regulatory network which act together to regulate and result in the enhancement of resistance to FOL in tomato.

The objective of this study was to determine differences between the transcript profiles of susceptible Moneymaker and resistant Motelle tomato cultivars in response to FOL infection and to reveal genes and miRNAs underlying the innate immune response against the fungal pathogen combining RNA-seq and sRNA-seq approach. In addition to genes predicted to response to pathogen infection, our results also uncovered a bunch of novel fungal pathogen-responsive miRNAs in tomato host plant for further functional characterization, and provided a broader view of the dynamics of tomato defense triggered by FOL infection.

## MATERIALS AND METHODS

### Tomato materials and fungal culture

For tomato materials and fungal culture, susceptible cultivar Moneymaker (*i-2*/*i-2*) and resistant cultivar Motelle (*I-2*/*I-2*) were applied for plant infection and libraries construction in this study. Tomato seedlings were grown at 25 °C with a 16/8-h light/dark cycle for two weeks used for all experiments. The wild-type FOL (race 2) strain is FGSC 9935 (known as FOL 4287 or NRRL 34936). Tomato seedlings were removed from soil and roots rinsed with tap water gently followed by incubating in the solution of FOL conidia at a concentration of 1×10^8^ conidia/ml for 30 minutes. Water treated tomato seedlings were used as the control. Forty seedlings were used for each treatment. All treated tomato seedlings were then replanted in soil and maintained in a growth chamber at 25 °C for 24 hours (16-hour light, 8-hour dark) as descripted previously (Robert-Seilaniantz *et al.* 2011). After infection, clean roots were collected and immediately frozen in liquid nitrogen for total RNA extraction. In order to minimize experimental variations, all root samples were consisted of pools of forty seedlings for each treatment. All experiments were repeated three times.

### RNA extraction, library preparation, and sequencing

To RNA extraction, library preparation, and sequencing, total RNA was isolated from roots using TRIzol^®^ Reagent (#15596026, Life Technologies, CA, USA) as descripted previously (Ouyang *et al.* 2014). For each sample, all roots from three biological repeats were pooled together for total RNA extraction.

For RNA-seq library construction, after the total RNA extraction followed by DNase I treatment, mRNA was enriched by magnetic beads with Oligo (dT). The mRNA was sheared into short fragments in the fragmentation buffer. Then cDNA was synthesized using the mRNA fragments as templates. cDNAs were purified and resolved with EB buffer for end reparation and single nucleotide A (adenine) addition followed by adding adapters to cDNAs. After agarose gel electrophoresis, the suitable cDNAs were selected for the PCR amplification as templates. During the quality control (QC) steps, Agilent 2100 Bioanaylzer and ABI StepOnePlus Real-Time PCR System were used in quantification and qualification of the sample library. The libraries were sequenced using Illumina HiSeqTM 2000.

For sRNA-seq library construction, 1 μg of total RNA were used for small RNA library generation. Briefly, total RNA samples were separated using polyacrylamide gel electrophoresis (PAGE), and cut out between 18 and 30 nt stripe to recover small RNA. 3’ and 5’ adapter were ligated at both ends, followed by reverse transcribed with Superscript II Reverse transcriptase using adapter-specific RT-primers. PCR products were then gel purified to enrich special fragments. The quality control (QC) steps were described as above. The purified high-quality cDNA library was sequenced using Illumina Genome HiSeq4000.

### Bioinformatics analysis of RNA-seq and sRNA-seq

Primary sequencing data that produced by Illumina HiSeqTM 2000/HiSeq4000, called as raw reads, were subjected to QC. After QC, raw reads were filtered into clean reads which were aligned to the reference sequences as described by previous report (Trapnell *et al.* 2012).

For RNA-seq, the clean reads were then aligned to the tomato reference genome downloaded from the Sol Genomics Network using Bowtie v0.12.5 (Langmead *et al.* 2009) and TopHat v2.0.0 (Trapnell *et al.* 2012; Trapnell *et al.* 2009) with default settings. Transcript abundance was calculated with Cufflinksv0.9.3 (Trapnell *et al.* 2010) based on fragments per kilo base of transcript permillion fragments mapped (FPKM) under default parameters settings. The transcript abundance was calculated for individual sample files followed further merged pairwise for each comparison (FOL inoculated between the two cultivars, and water treated between the two cultivars and FOL inoculated versus water treated for each cultivar) using Cufflinks utility program-Cuffmerge (Trapnell *et al.* 2012). The pairwise comparisons of gene expression profiles between the two populations were done using the Cuffdiff program of the Cufflinks version 1.3.0 (Trapnell *et al.* 2010). The genes were considered significantly differentially expressed if Log_2_ FPKM (fold change) was ≥1.0 and false discovery rate (FDR, the adjusted P value) was <0.01. The q-value which was a positive FDR analogue of the p-value was set to <0.01 (Storey *et al.* 2003).

For sRNA-seq, the sequence data were subsequently processed using in-house software tool SeqQC V2.2. House-keeping small RNAs including tRNAs, rRNAs, snoRNAs and snRNAs were removed by blasting in GenBank (http://www.ncbi.nih.gov/Genbank) servers. The trimmed reads were then aligned to the tomato reference genome downloaded from http://www.mirbase.org (miRBase 21.0) and http://www.ncbi.nlm.nih.gov using Bowtie v0.12.5 (Oliveros *et al.* 2007-2015) and TopHat v2.0.0 (Trapnell *et al.* 2009; Ruepp *et al.* 2004) with default settings. To identify known miRNAs, the remaining unique small RNA sequences were then aligned against the miRBase 21.0 allowing maximum one mismatch. After assigning the known miRNA sequences into their respective groups or families, rest of the sequences were checked for novel miRNAs.

### Functional categorization of DEGs

DEGs were functionally categorized online for all pairwise comparisons according to the Munich Information Center for Protein Sequences (MIPS) functional catalogue (Ruepp *et al.* 2004). The functional categories and subcategories were regarded as enriched in the genome if an enrichment p-value was below <0.05. The Kyoto Encyclopedia of Genes and Genomes (KEGG) pathway analyses were performed using interface on blast2GO (Blast2GO v2.6.0, http://www.blast2go.com/b2ghome) for all DEGs to identify gene enrichment on a specific pathway.

### Gene Ontology (GO) and pathway enrichment analysis

Gene Ontology (GO) and pathway enrichment were performed using DAVID software (Smyth *et al.* 2005). Graphs of the top 20 enriched GO terms for each library were generated using the Cytoscape Enrichment Map plugin (Smoot *et al.* 2005; Merico et al. 2010).

### Quantitative RT-PCR (qRT-PCR) and Northern blot analysis

Quantitative RT-PCR (qRT-PCR) and Northern blot analysis were performed according to our previous protocol [32]. Briefly again, expression of DEGs were determined using quantitative RT-PCR (qRT-PCR). cDNAs were generated from 1 μg of total RNA using the SMART MMLV Reverse Transcriptase (Takara, Mountain View, CA) followed by diluting two times and using as template for quantitative RT-PCR, which was performed with the CFX96 real-time PCR system (Bio-Rad, Hercules, California, USA). Primers used for qRT-PCR were designed from 3-UTR for individual gene. Each reaction mixture (20 μL) contained 1 μL of cDNA template, 10 μL of SYBR1 Green PCR Master Mix (Applied Biosystems, Foster, CA) and 1 μL of each primer (10 μM). For each cDNA sample, three replications were performed. The expression level of 18S rRNA was used as internal control for normalize the expression level of selected genes, and were calculated as the fold change by comparison between in FOL treated and water in treated samples.

Tomato root total RNA were used for small RNA northern blot analysis, For each sample, 40 μg total RNA was resolved on urea denaturing polyacrylamide gels (Urea-PAGE). miRNA-specific oligonucleotide probes were end-labeled using γ-^32^P-ATP (#M0201, New England Biolabs, Ipswich, MA). Gel staining with ethidium bromide was used as a loading control. All blots were imaged by PhosphorImager (GE Life Sciences, Pittsburgh, PA, USA). The images were cropped and adjusted with brightness and contrast in Photoshop CS6 from original digital images.

### Statistical analyses

All data in this study were analyzed with ANOVA program or Student’s t-test analysis using SPSS 11.5 (SPSS Company, Chicago, IL) for statistical analysis.

## RESULTS

We recruited tomato cultivars susceptible Moneymaker (*i-2/i-2*) and resistant Motelle (*I-2/I-2*) genotypes. Motelle plants displayed strong resistance to FOL infection. Four weeks after FOL infection, Moneymaker plants exhibited severe wilting symptoms (Fig. 1A). We generated four libraries for RNA-seq and sRNA-seq, respectively, including Moneymaker treated with FOL/water (MM_FOL/MM_H_2_O), Motelle treated with FOL/water (Mot_FOL/Mot_H_2_O). Therefore, totally eight libraries were constructed. After treating with FOL/water, total RNA from tomato roots were extracted for both RNA-seq and sRNA-seq. By taking advantage of these two cultivars, we combined RNA-seq and sRNA-seq to explore the responsive crucial molecular players and pathways. A briefly mRNA and miRNA detection workflow was presented to summarize the method utilized in this study (Fig. 1B).

**Figure 1.**
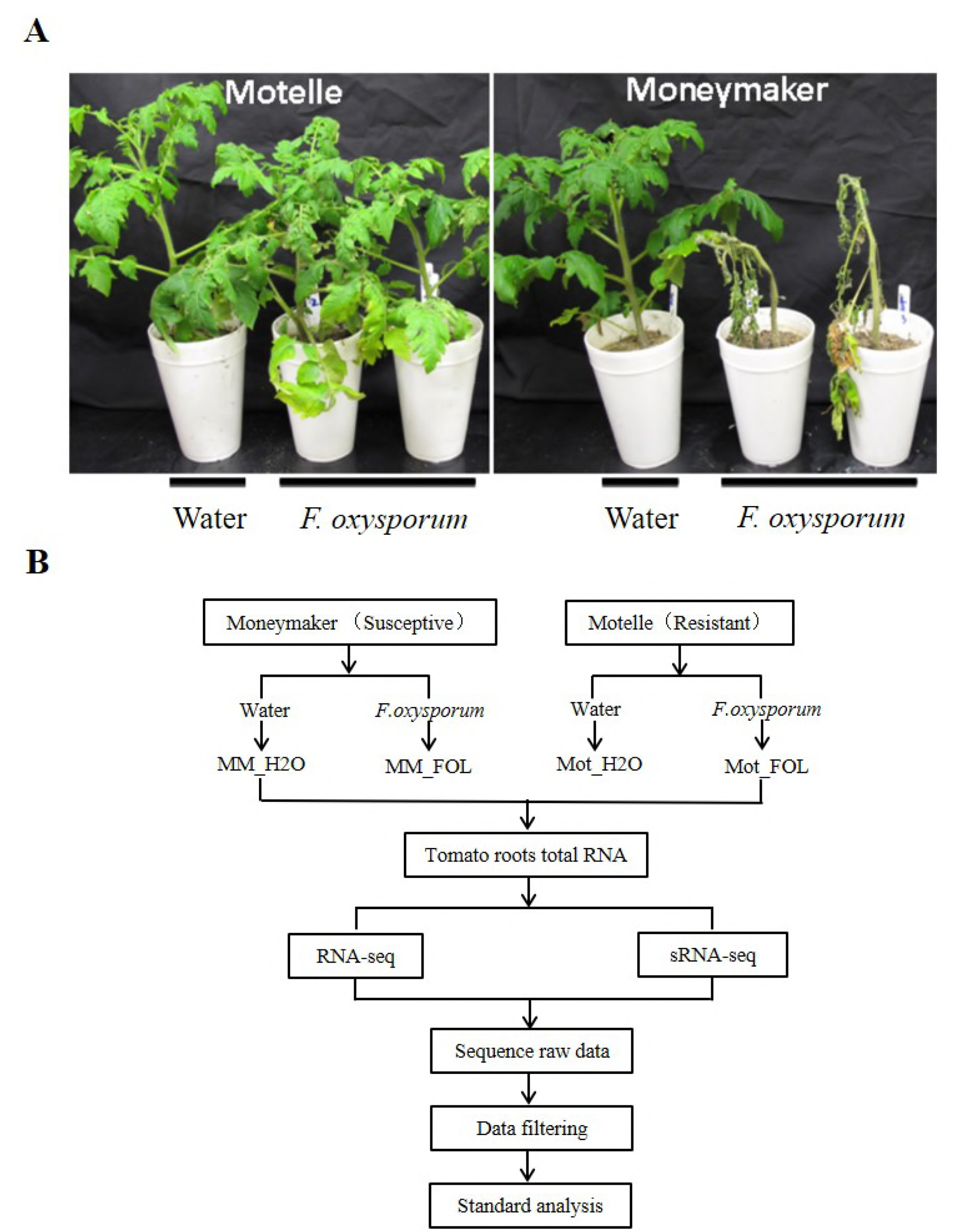
Post inoculation phenotype in susceptible cultivar Moneymaker and resistant cultivar Motelle. A The phenotype of tomato seedlings infected by FOL. Two-week-old tomato seedlings were treated with FOL or water followed by photographing four weeks later. B Briefly mRNA and miRNA detection workflow.

Total RNA from three independent sets of plants (biological replicates) was extracted from roots. RNA-seq libraries were developed from four treatments. Each RNA-seq library included three biological replicates. Using Illumina sequencing, after quality control filtering and trimming adaptor sequences, a total of more than 182.6 million high quality clean reads were collected by RNA-seq from the four libraries. The sequencing yielded ~ 46 million clean reads from each library respectively. The clean reads were then mapped to the *Solanum lycopersicum* genome using the Tomato Genome assembly (SL3.0) and annotations (ITAG3.0) (www.solgenomics.net/). Approximately three fourths of the total Illumina reads perfectly matched the genome or gene and were used for further analysis. Analysis of the expressed transcripts in the libraries showed that almost equal number of transcripts were observed by ~23 million in the four libraries (Table 1). We used Cufflink to measure the expression level of Tomato annotated genes. Among these transcripts, 21,189 (86.4%) genes were expressed in all four libraries, and 382 (1.6%), 333 (1.4%), 357 (1.5%) and 372 (1.5%) genes were uniquely presented in MM_H_2_O, MM_FOL, Mot_H_2_O and Mot_FOL library, respectively (Fig. 2A, Table S1).

**Figure 2.**
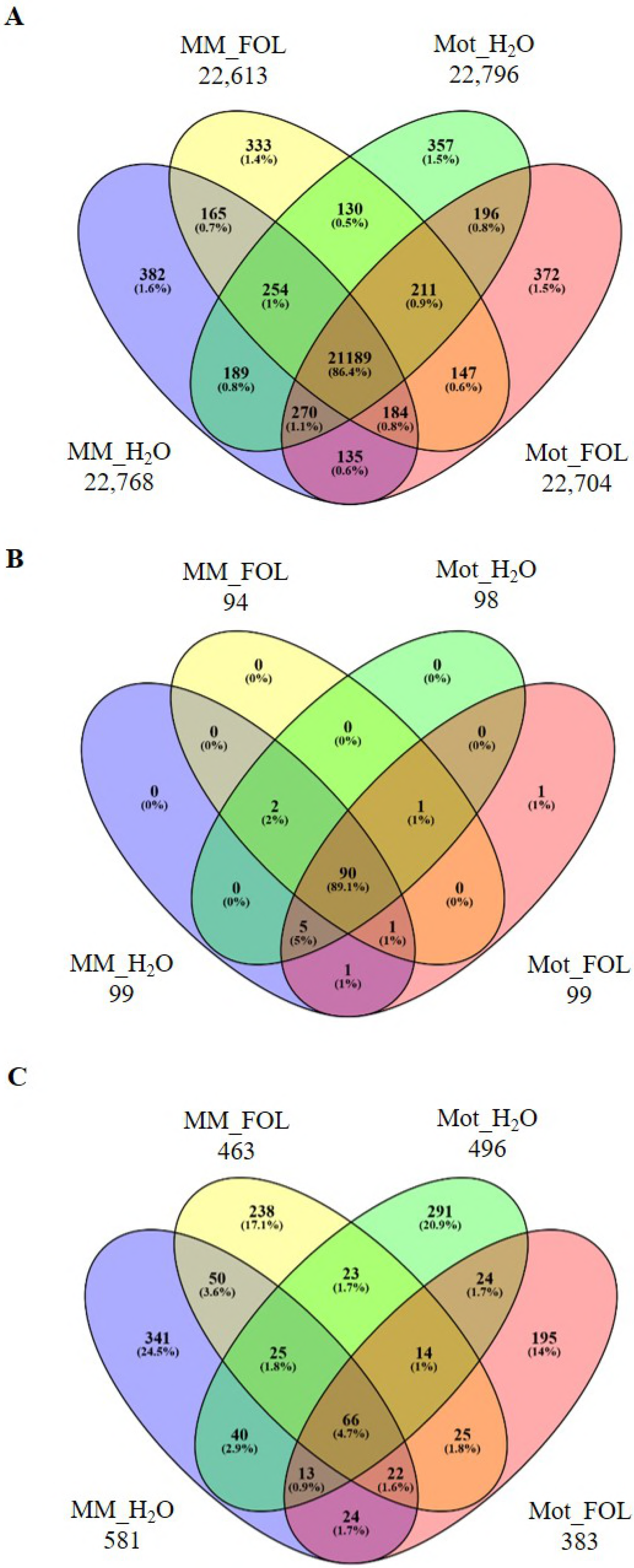
The Venny diagrams showing the overlaps among four comparisons of FOL and water treatment. A mRNAs. B Known miRNAs. C Novel miRNAs.

**Table 1.** Summary of RNA-seq and sRNA-seq datasets from four libraries.

|  | Annotation | MM_H<sub>2</sub>O | MM_FOL | Mot_H<sub>2</sub>O | Mot_FOL |
| --- | --- | --- | --- | --- | --- |
| RNA-seq | Clean reads | 45,616,330 | 45,635,428 | 45,680,034 | 45,661,734 |
|  | Genome map rate | 75.49% | 67.89% | 75.87% | 70.46% |
|  | Gene map rate | 76.87% | 68.47% | 75.92% | 71.26% |
|  | Expressed gene | 22,796 | 22,639 | 22,825 | 22,725 |
|  | Novel gene | 775 | 761 | 795 | 705 |
|  | Alternative splicing | 32,482 | 32,689 | 33,706 | 32,965 |
|  | Total reads apped to genome | 75.49% | 67.89% | 75.87% | 70.46% |
|  | Perfect match to genome | 63.00% | 55.52% | 62.61% | 57.41% |
|  | Mismatch to genome | 12.49% | 12.38% | 13.26% | 13.05% |
|  | Unique match to genome | 74.32% | 66.85% | 74.79% | 69.46% |
|  | Total Unmapped reads | 24.51% | 32.11% | 24.13% | 29.54% |
| sRNA-seq | Clean reads | 31,192,441 | 34,969,928 | 33,000,743 | 33,125,499 |
|  | snRNA | 233,962 | 413,635 | 360,602 | 416,831 |
|  | rRNA | 15,246,656 | 16,245,295 | 18,695,564 | 17,611,875 |
|  | snoRNA | 88,371 | 112,863 | 90,081 | 138,206 |
|  | Repeat | 52,8140 | 318,782 | 427,681 | 310,377 |
|  | miRNA | 339,025 | 165,398 | 219,845 | 143,398 |
|  | tRNA | 655,213 | 715,878 | 642,906 | 712,388 |

By sRNA-seq, we collected a total of more than 132.2 million high quality clean reads including 31,192,441 from MM_H_2_O, 34,969,928 from MM_FOL, 33,000,743 from Mot_H_2_O, and 33,125,499 from Mot_FOL, respectively. Valid sequences were classified according to the genomic regions matched. The composition of sRNA pool was comprehensive and contained a huge portion of other non-coding RNA species including snRNA, rRNA, snoRNA, repeat, miRNA and tRNA. Of these sRNA, we detected 339,025 miRNAs from MM_H_2_O, 165,398 from MM_FOL, 219,845 from Mot_H_2_O, and 143,398 from Mot_FOL (Table 1, S1). Synchronously, a venny diagram was recruited to demonstrate the distribution of miRNAs among the four comparisons. To known miRNAs, we could see that the majority of known miRNAs were overlapped among these four libraries. To novel miRNAs, however, there were 341 (24.5%), 238 (17.1%), 291 (20.9%) and 195 (14%) miRNAs altered the expression uniquely in MM_H_2_O, MM_FOL, Mot_H_2_O and Mot_FOL samples, respectively (Fig. 2B).

Differentially expressed genes (DEGs) were defined as genes with fold-change >2 folds and FDR < 0.01. A total number of 3,942 and 4,168 genes were showed significantly differential expression in MM_FOL vs. MM_H_2_O library and Mot_FOL vs. Mot_H2O library, respectively. A majority of these DEGs were overlapped in both water and FOL treated two tomato cultivars. Among these DEGs, 221 were down-regulated in MM_FOL vs. MM_H_2_O and 219 were down-regulated in Mot_FOL vs. Mot_H_2_O, while 261 were up-regulated in MM_FOL vs. MM_H_2_O and 415 up-regulated in Mot_FOL vs. Mot_H_2_O (Fig. 3, detailed in Table S1).

**Figure 3.**
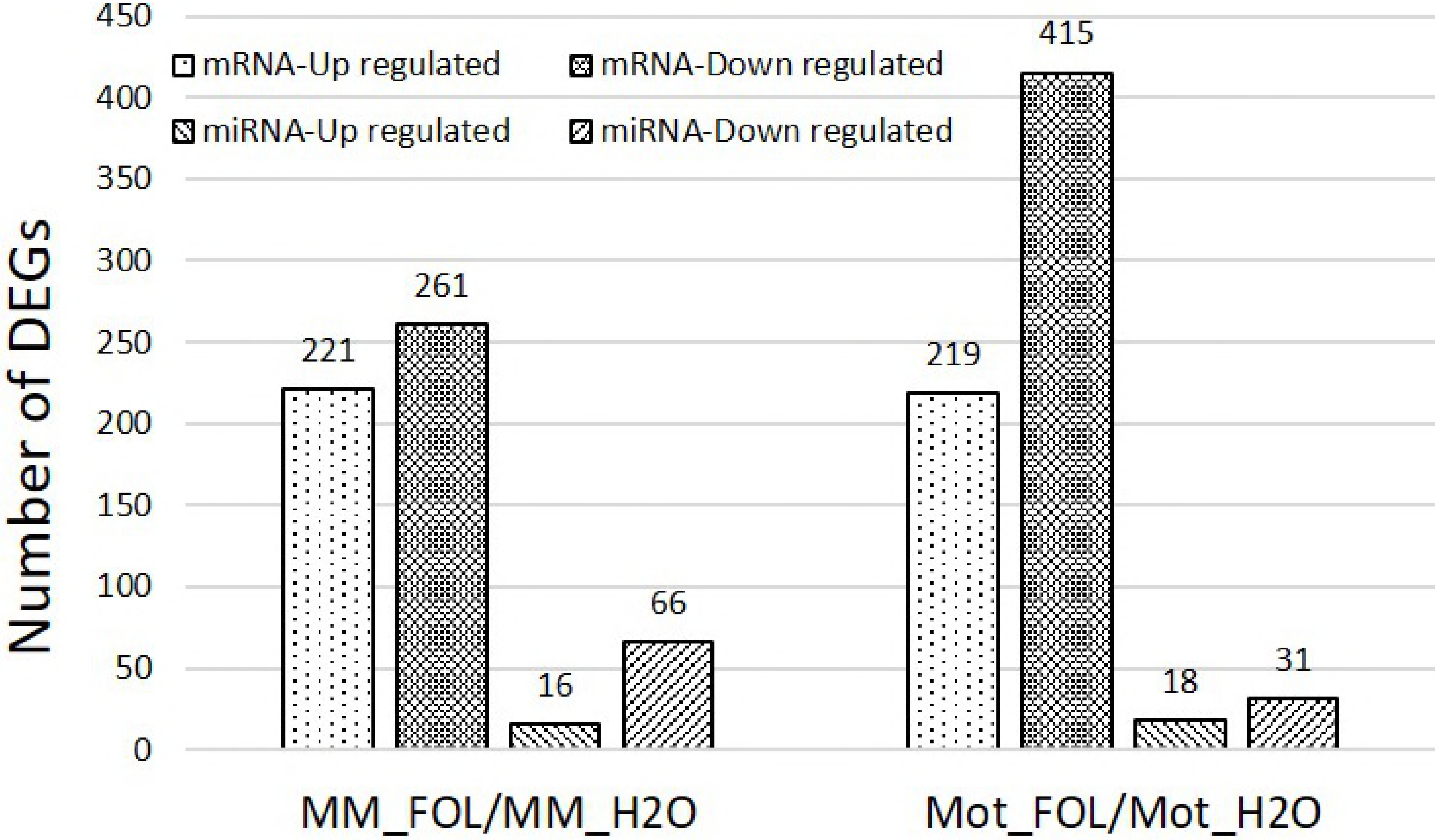
Statistics of Differentially Expressed mRNAs and miRNAs between FOL and water treatment.

Screening of DE miRNAs, the results displayed that the miRNA expression level changed under FOL treatment. The histogram was shown that fewer miRNAs have altered expression after FOL treatment in Motelle sample (18 up-regulated and 31 down-regulated) compared to in Moneymaker sample (16 up-regulated and 66 down-regulated) (Fig. 3). Taken together, FOL treatment had a significant impact on global gene/miRNA expression profile in tomato plants.

To explore the distribution of DEGs/DE miRNAs, gene ontology (GO) enrichment analyses were conducted based on these DEGs to elucidate the biological processes/pathways. A total of 530 and 769 GO terms were discovered in MM_FOL vs. MM_H_2_O and Mot_FOL vs. Mot_H_2_O library, respectively, related to three main classes: Biological Processes, Cellular Component, and Molecular Function. GO enrichment analysis revealed that GO terms were mainly classified into catalytic activity (104 out of 530 in MM_FOL vs. MM_H_2_O library, and 141 out of 769 in Mot_FOL vs. Mot_H_2_O library) (the same define in the following text), metabolic process (81 out of 530, and 118 out of 769), and binding (72 out of 530, and 104 out of 769). For the class of response to stimulus, however, no significant change was presented between these two libraries (31 out of 530, and 36 out of 769) (Fig. 4A, Table S2). To DE miRNAs, cellular process and metabolic process were the two most represented categories in Biological Process, being associated with 23.8% of the coding regions of Moneymaker, and 20.8% of the coding regions of Motelle. The cell and cell part category were associated with 21.7% of the transcripts in Moneymaker, and 22.6% in Motelle, being the two most represented in the Cellular Component class. Majority of the annotated transcripts were found to be associated with the binding category and catalytic activity in Molecular Function, with 19.9% of the transcripts in Moneymaker and 20.8% in Motelle (Fig. 4B, Table S3). It was worthy to note that when generally compared the two deep sequencing results (RNA-seq and sRNA-seq), we found that there were more regulated transcripts responding to FOL invasion in Motelle than that of in Moneymaker by RNA-seq, on the contrary, more regulated miRNAs responding to FOL invasion in Moneymaker than that of in Motelle by sRNA-seq.

**Figure 4.**
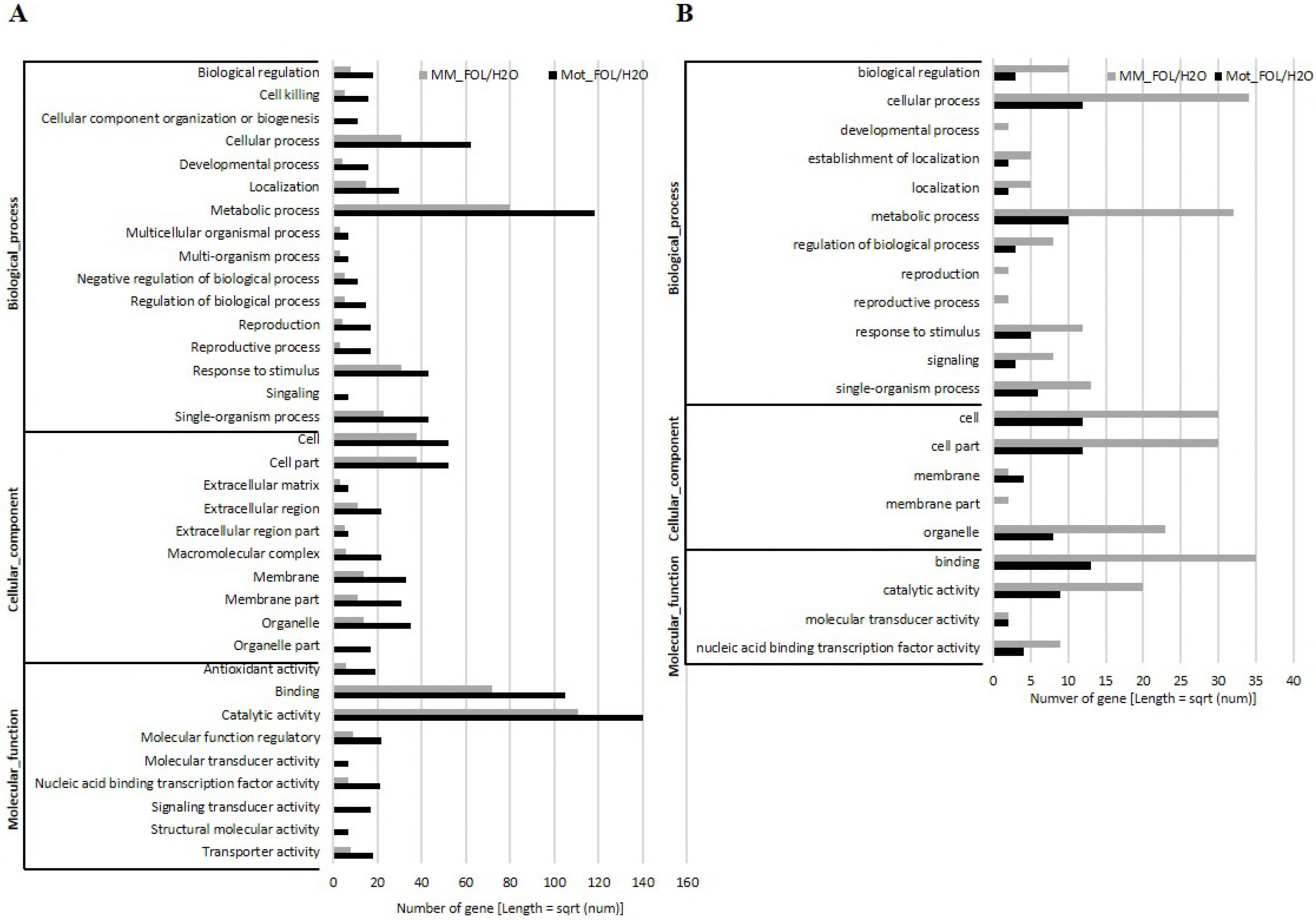
Functional categorization of significantly Differentially Expressed mRNA and miRNA under FOL invasion in tomato. The results were basically summarized into three main categories: biological processes, cellular components, and molecular functions. All statistically significant genes from four libraries were assigned to GO terms. A mRNA from RNA-seq. B Targets of miRNAs from sRNA-seq.

To further understand the biological functions, pathway enrichment of DEGs were performed to discover the effect of FOL to host plant. Be worth mentioning, plant-pathogen pathway was ranked in the 29^th^ (24 out of 356 DEGs) in MM_FOL vs. MM_H_2_O group (Detailed in Table S4), however, it was presented in the 8^th^ (40 out of 469 DEGs) in Mot_FOL vs. Mot_H_2_O group (Detailed in Table S5). These results may indicate that more pathogen resistance genes are regulated in resistant cultivar Motelle than that in susceptible cultivar Moneymaker. To summarize, these DEGs included genes encoding WRKY protein (8 genes), receptor kinase (20 genes), MYB transcription factor (7 genes), NBS-ARC protein (18 genes), Calmodulin-like protein (9 genes), MAPK (2 genes) and others (7 genes). Seventeen DEGs were regulated in both Moneymaker and Motelle. (Table 2).

**Table 2.**
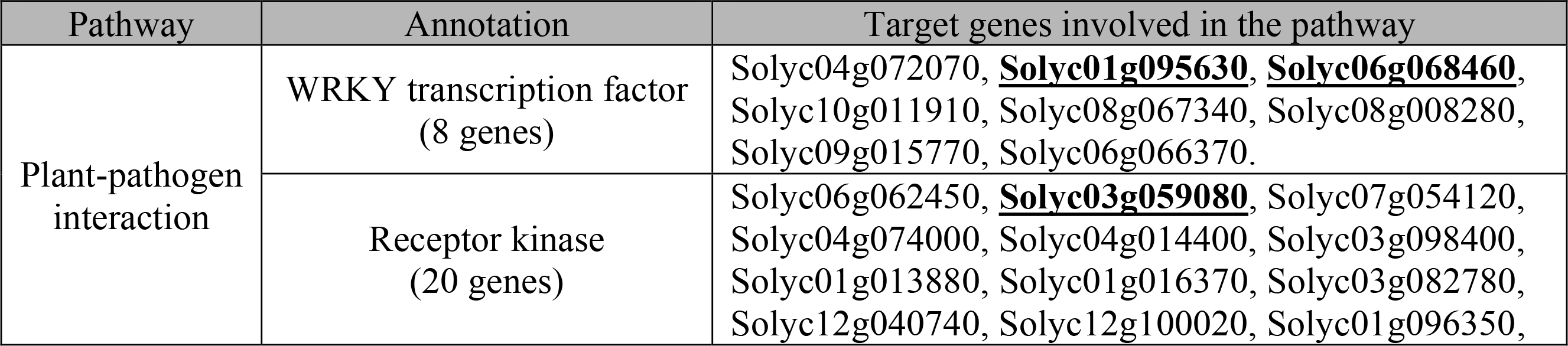
Detail information of regulated genes involved in the plant-pathogen interaction pathway. Genes (bold) were regulated in both Moneymaker and Motelle. Genes (underlined) were further analyzed by qRT-PCR.

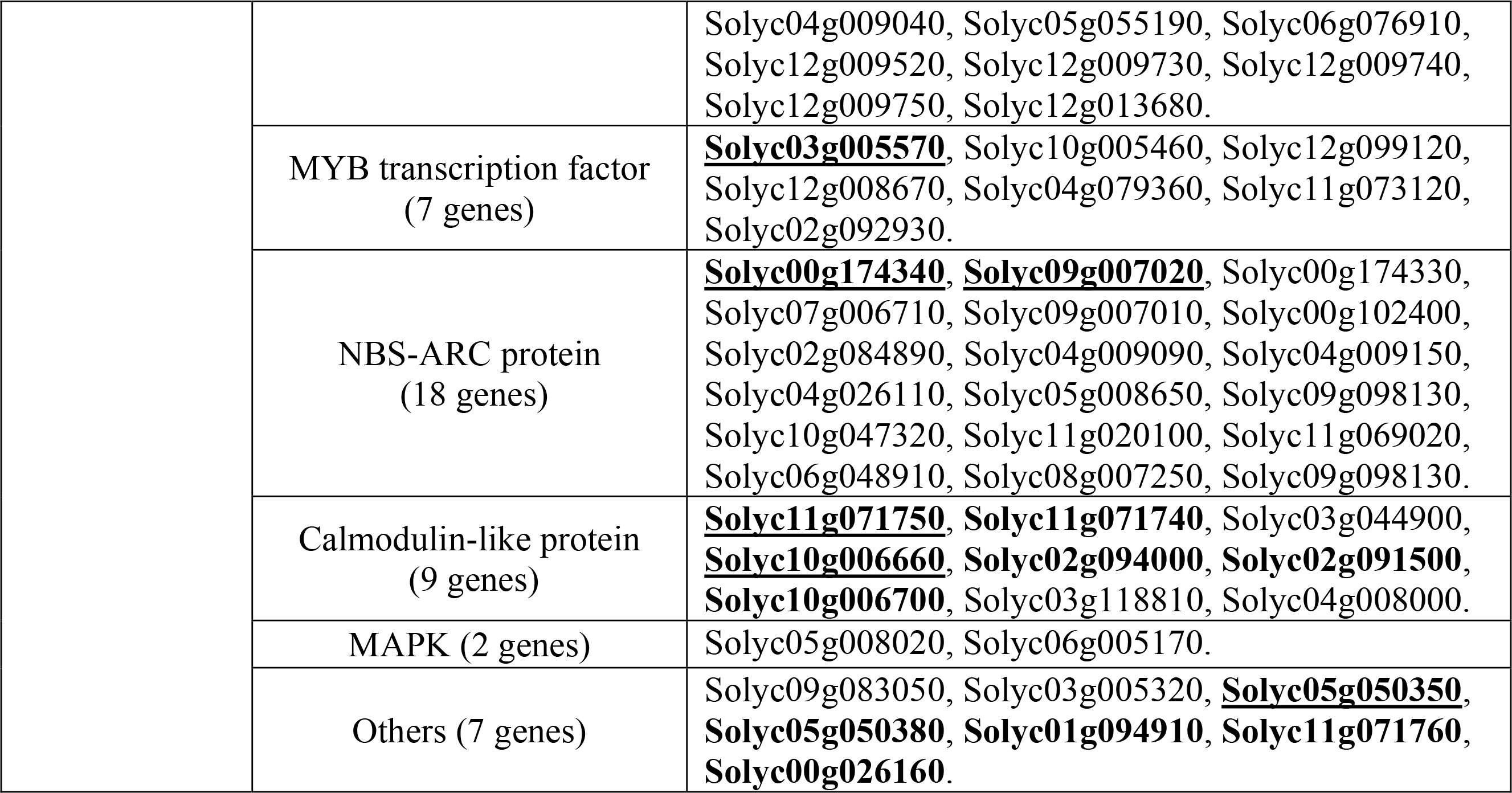

To miRNAs from sRNA-seq, we predicted the targets of known and novel miRNAs with the number 1179 and 2615, respectively (Table S6). In total, thirty-two miRNA families were detected in all libraries in both Moneymaker and Motelle under FOL infection (Detailed in Table S7). Of these miRNA families, eleven miRNA families were expressed specifically in *Solanaceae* plants. Among these specific expressed miRNA families, five of them, including miR6022, miR6023, miR6026, miR6027 and miR6024, were predicted regulating plant innate immune receptors which were listed in Table 3.

**Table 3.**
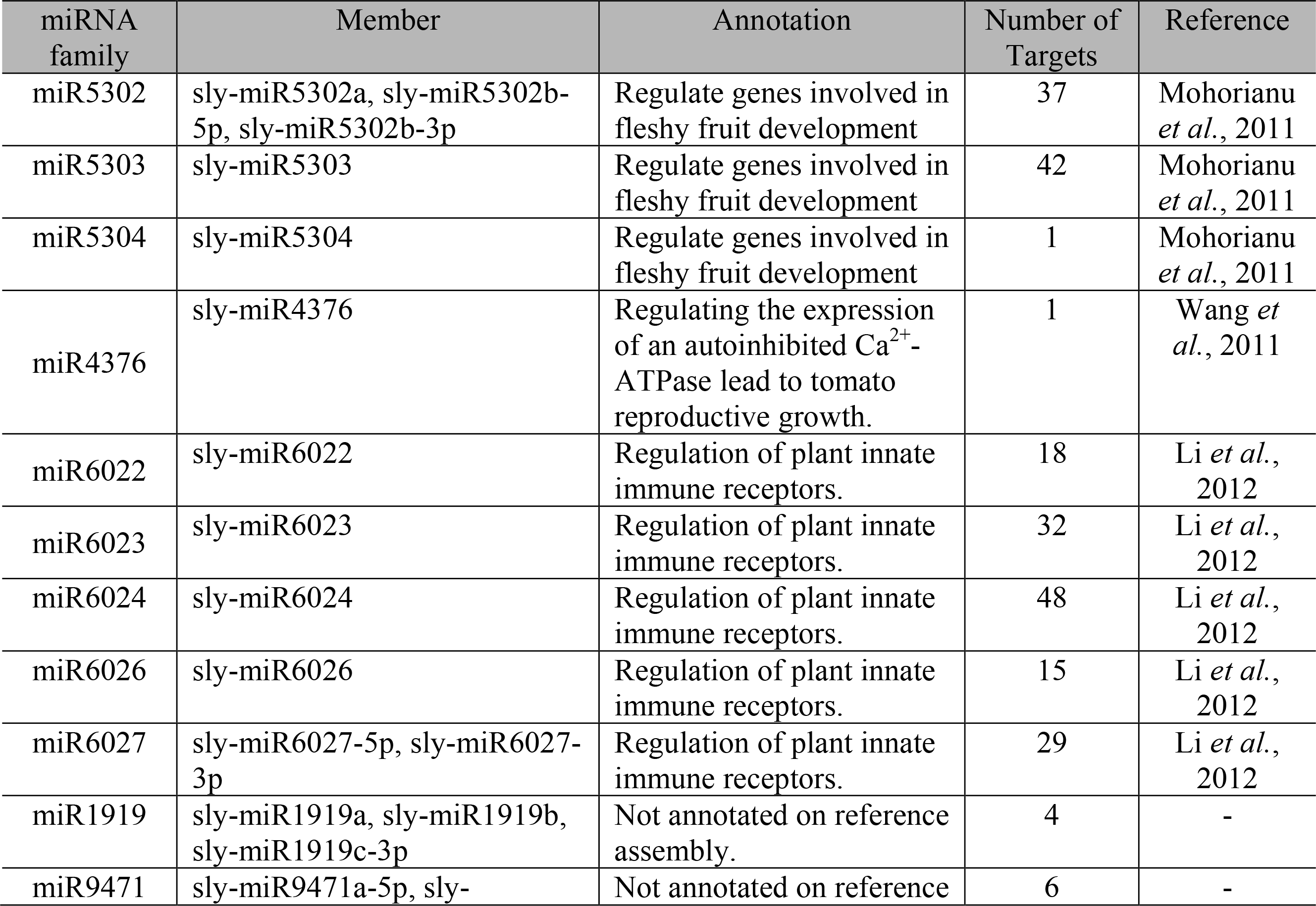
Function description of miRNA families especially presented in the nightshade family (*Solanaceae* plant).

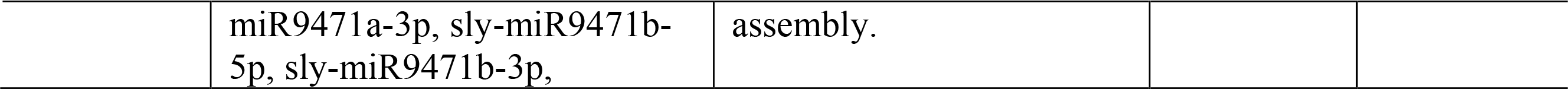

To verify the DEGs in plant-pathogen interaction pathway, nine predicted disease related DEGs, regulated in both Moneymaker and Motelle, were selected to characterize the gene expression profiles between FOL and water treated Moneymaker/Motelle by qRT-PCR using primers listed in Table S8. These DEGs included Solyc01g095630 (SlWRKY41), Solyc06g068460 (SlWRKY40), Solyc03g059080 (Receptor-like serine/threonine protein kinase), Solyc03g005570 (Myb-related transcription factor), Solyc00g174340 (Pathogenesis-related protein 1b), Solyc09g007020 (Pathogenesis-related protein), Solyc11g071750 (Calmodulin-like protein), Solyc10g006660 (Calcium-binding EF hand family protein) and Solyc05g050350 (Cyclic nucleotide gated channel). The results of qRT-PCR showed the similar pattern to sequencing results with minute difference. Among these DEGs, Solyc01g095630, Solyc03g059080, Solyc00g174340, Solyc11g071750 and Solyc05g050350 were induced greatly in resistant cultivar Motelle affected by FOL, however, no significant changes were presented in susceptible cultivar Moneymaker between FOL and water treatment. Solyc06g068460 was induced in both Moneymaker and Motelle upon FOL treatment. On the other hand, Solyc11g071750 and Solyc10g006660 were suppressed in both Moneymaker and Motelle plants when treated with FOL (Fig.5)

**Figure 5.**
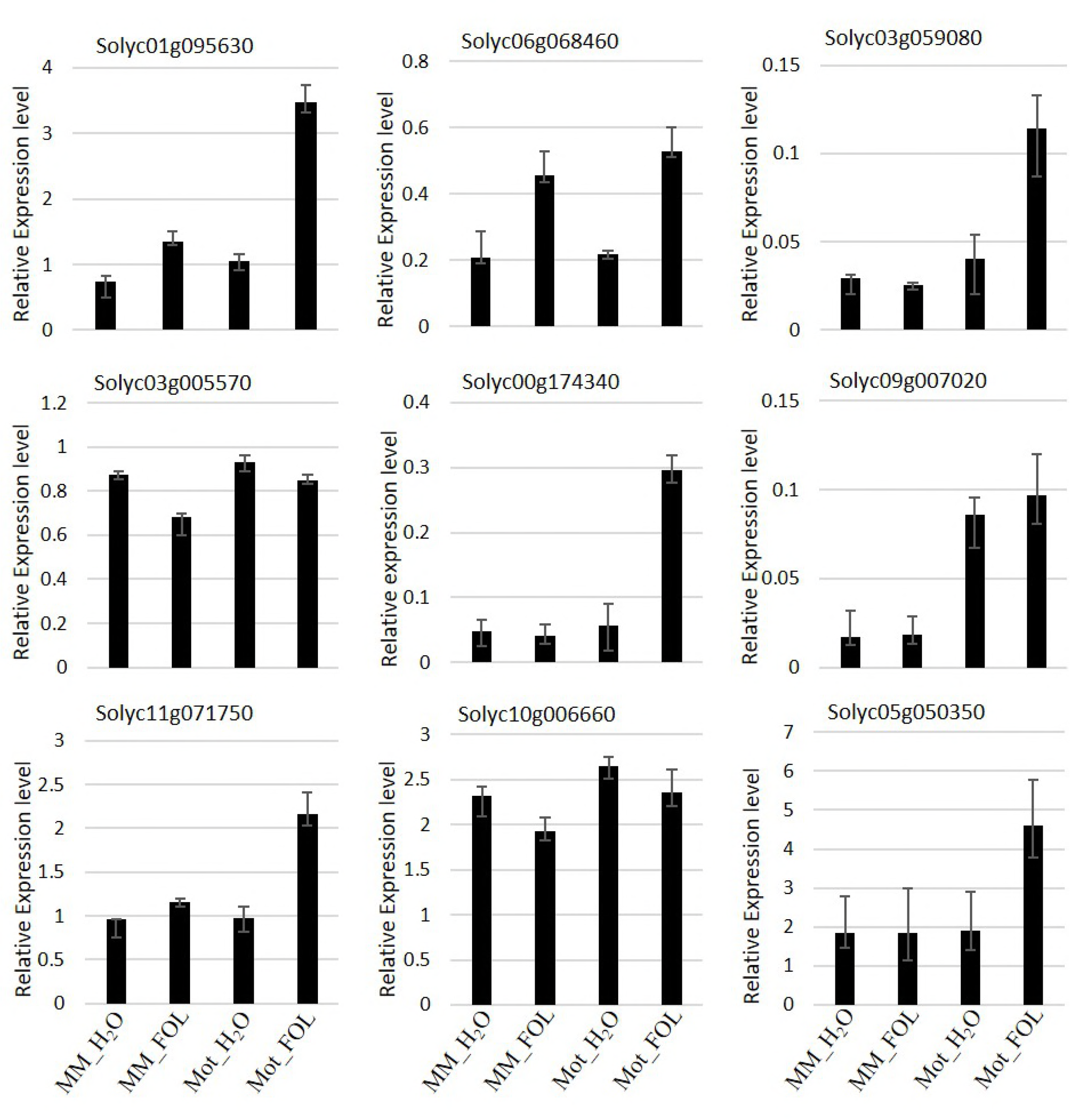
Validation of Differentially Expressed genes selected in plant-pathogen interaction pathway by qRT-PCR. Total tomato root RNA was reverse transcribed to cDNA used as template for qRT-PCR with gene-specific primers. Each column represents an average of three replicates, and error bars represent the standard error of means.

We also selected additional twenty defense-related genes at the same level of sample read coverage to further verify the data from RNA-seq using qRT-PCR. The results displayed the same expression of up-regulation or down-regulation but with the fold changes varies due to the two different methods for gene quantification (data not presented).

To further characterize the miRNAs expression patterns, we identified twenty-two miRNAs up or down-regulated in both Moneymaker and Motelle under FOL/water treatment (Fig. 6). To test the reliability of our sRNA-Seq data, Northern blot analysis was performed on nine DE miRNAs (sly-miR160a, sly-miR477-5p, sly-miR167a, novel_mir_273, novel_mir_469, novel_mir_365, novel_mir_675, novel_mir_504 and novel_mir_762) using primers listed in Table S8. Ethidium bromide staining of gels were applied as loading control. Our data indicated that Sly-miR477-5p, sly-miR167a, novel_mir_675, novel_mir_504 and novel_mir_762 were repressed congruously in both Moneymaker and Motelle when treated with FOL. Novel_mir_365 and novel_mir_469 were slightly up-regulated in Motelle under FOL invasion (Fig. 7). These data displayed a credible consistence with the results of sRNA-seq, showing the reliability of sRNA-seq analysis.

**Figure 6.**
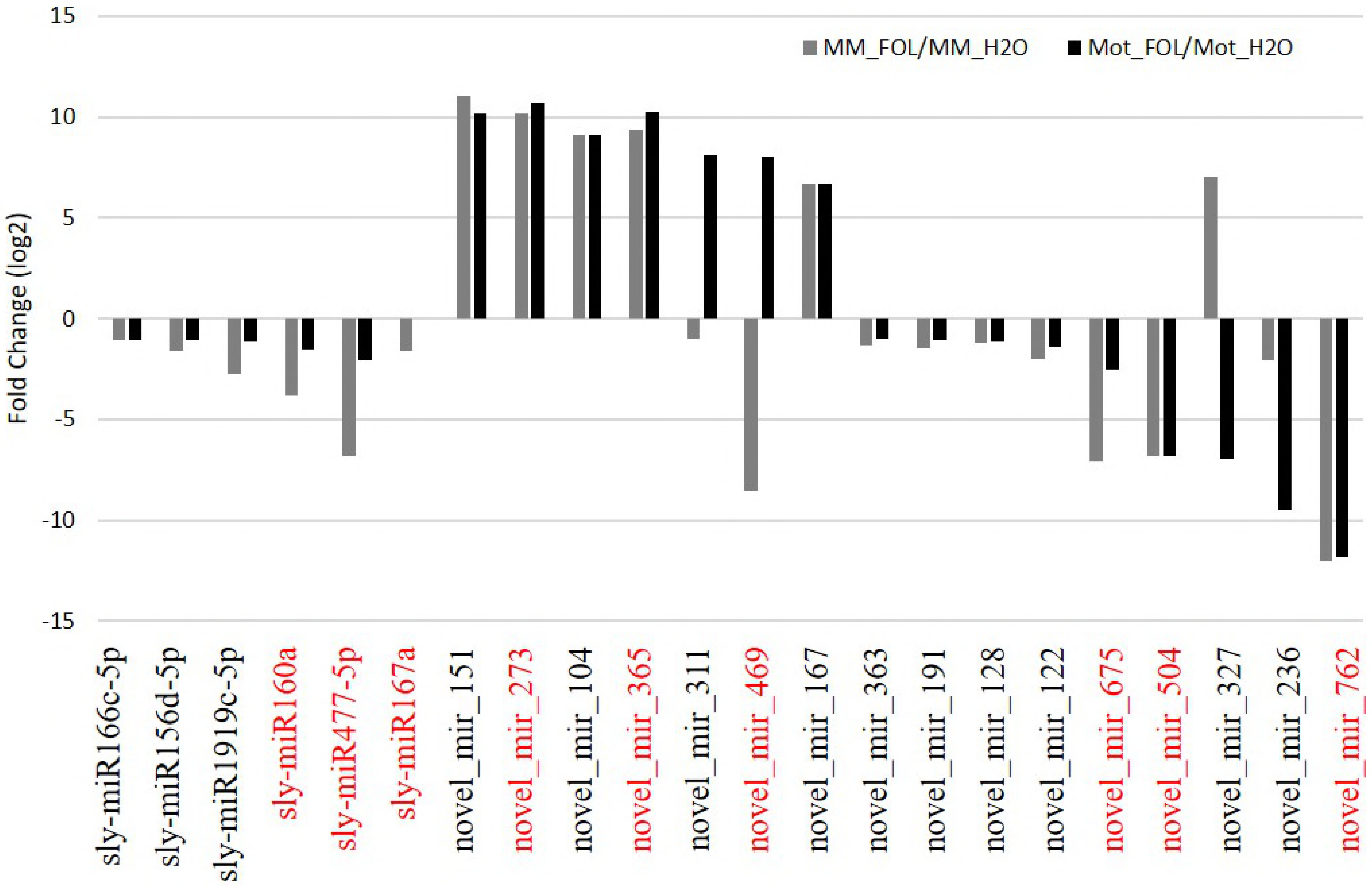
Profiling of miRNAs response to FOL in tomato plants. According to sRNA-seq analysis, partial of regulated miRNAs were summarized by normalizing reads of water treatment for each cultivar.

**Figure 7.**
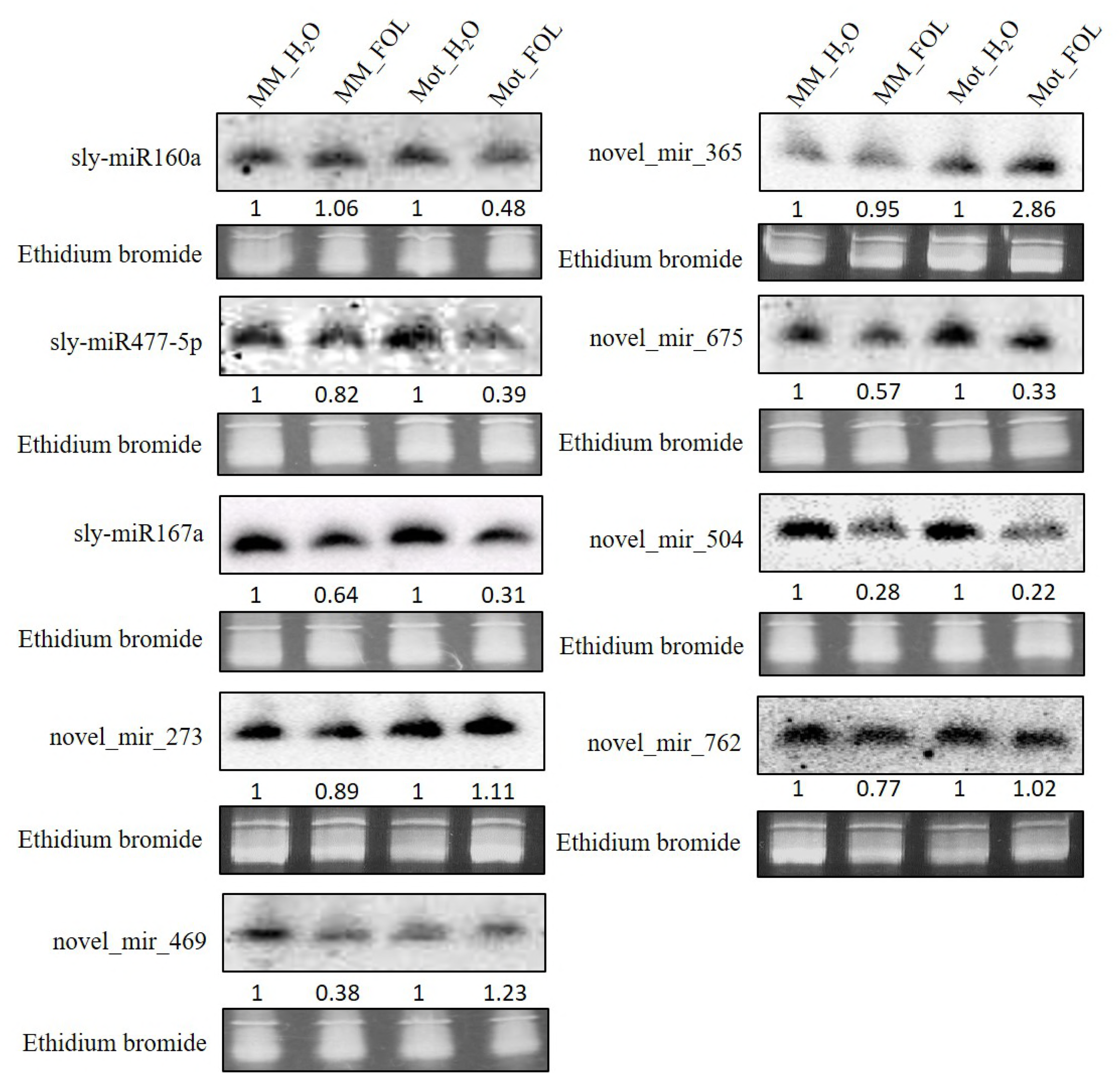
Expression validation of selected miRNAs by Northern blot analysis. Several miRNAs including known and novel highlighted with red in Fig. 6 were selected randomly for Northern blot analysis. Total root RNA samples (40 μg) were from four treatments. Gel staining with ethidium bromide were used as loading control for each blot. Blots were imaged using a Phosphorimager. Using ImageJ software to measure the grey density, the numbers below each blot present the relative enrichment of individual miRNA in each treatment normalized to the corresponding water-treated control.

## DISCUSSION

In the present study, we explored the advantages of near-isogenic susceptible and resistant cultivars of tomato infected by FOL to uncover a global mRNA and miRNA expression profile of tomato-FOL interaction using Illumina sequencing, which helped us dissect the molecular mechanisms underlying FOL invasion. The components of plant response to pathogen challenging may lead to understand the potential defense mechanisms. Plants have evolved a complicate defense system against pathogens including cascade signaling activation, the regulation of gene expression, synthesis of defensive metabolites as well as hormone balancing (Mukhtar *et al.* 2011; Andolfo *et al.* 2014). So far, by taking advantage of high-throughput RNA sequencing (RNA-seq) approach, a few of transcriptome studies discovering the FOL-host interaction have been reported in plants such as banana, watermelon, mango and *Arabidopsis* (Guo *et al.* 2014; Xing *et al.* 2016; Liu *et al.* 2015; Liu *et al.* 2016; Chen *et al.* 2014; Gupta *et al.* 2014), shedding light on the cross-talking among different signaling pathways involving in plant-pathogen interaction.

When plant is attacked by pathogen, the host reprograms metabolism balance between development and the resources to support defense to pathogen, involving biological process, cellular components and molecular functions (Mithöfer *et al.* 2012). Based on our results, the tomato-FOL interaction basically follows the typical reaction of biotrophic phase pathogens infection. Gene Ontology analysis of DEGs between two tomato cultivars reveals specific enriched categories in both interactions. In resistant tomato cultivar Motelle, cellular component organization or biogenesis, signaling, molecular transducer activity, and signal transducer activity were evidenced when compared to susceptive tomato cultivar Moneymaker. Among them, cellular component organization and biogenesis are critical metabolic activities required by plants to survive under fungus-inflicted stresses (Paul *et al.* 2011). Generally, the genes involved in GO analysis present in Motelle more than in Moneymaker upon FOL infection which may due to different resistant cultivar.

Two main mechanisms, pathogen-associated molecular patterns (PAMPs) or named as microbe-associated molecular patterns (MAMPs) (Boller *et al.* 2009; Cui *et al.* 2014; Yang *et al.* 2014) and the adaptive immune system composed of resistant (R) genes (Dangl *et al.* 2011; Van Ooijen *et al.* 2007; Marone *et al.* 2013), are involved in plant responses to pathogenic microorganisms in plant. At least five different classes of *R* genes have been classified based on functional domain (Marone *et al.* 2013). Among these classes, a nucleotide-binding site (NBS) and leucine-rich repeats (LRRs) (NBS-LRR) is known as the most numerous R-gene class (Dangl *et al.* 2011). Previously, we reported that tomato endogenic miRNA slmiR482f and slmiR5300 conferred tomato wilt disease resistance. For each of slmiR482f and slmiR5300, two target genes were predicted encoding protein with full or partial NBS domains respectively, confirmed to exhibit function of resistance to FOL (Ouyang *et al.* 2014). A few of investigations have been demonstrated that NBS-LRR proteins recognize a specific Avr and display disease resistance in several plant species, including rice, tomato, *N*. *benthamiana*, *Arabidopsis* and wheat Ouyang *et al.* 2014; Sinapidou *et al.* 2004; Peart *et al.* 2005; Lee *et al.* 2009; Loutre *et al.* 2009; Narusaka *et al.* 2009; Okuyama *et al.* 2011). Transcriptional regulation of defense genes has been known as a central in plant defense responses. Certain a few of plant TF families, such as AP2/ERF, bHLH, TGA/bZIP, MYB, NAC and WRKY, appear to be prominent regulators of host defense (Tsuda *et al.* 2015). Our results also revealed that WRKY and MYB transcription factors were involved in the response to FOL invasion in tomato. Comprehensive studies have firmly established that WRKY TFs cope with divergences biological processes but most functionally required for plant immunity in tomato, *Arabidopsis*, barley and rice (Birkenbihl *et al.* 2017). Several MYB proteins, including AtMYB30, AtMYB44, AtMYB108/BOSI1 and HvMYB6, demonstrate prominent functions in plant immunity (Buscaill *et al.* 2014). However, there was no reported MYB protein conferring disease resistance in tomato species, so far.

It is well-established that miRNA is one of the plant produced two major classes of endogenous small RNAs, mediating sequence-dependent post-transcriptional gene silencing (PTGS) by guiding mRNA cleavage and/or translation inhibition. In the past decades, genome-wide small RNA analyses using sRNA-seq approach have been conducted for several plant-filamentous pathogen interactions (Ouyang *et al.* 2014; Inal *et al.* 2014; Radwan *et al.* 2011; Baldrich *et al.* 2014; Shapulatoy *et al.* 2016). However, so far, only miR482/2118 superfamily has been elucidated for the disease resistance function in tomato (Ouyang *et al.* 2014; Shivaprasad *et al.* 2012). From our sRNA-seq data, a common set of plant innate immunity miRNAs were regulated after FOL infection in both susceptible and resistant tomato plant. Not surprised, more miRNAs were presented to respond to FOL invasion in Motelle than that of in Moneymaker. Further potential target prediction results by online tool psRNATarget indicated that a few of these targets were pathogen resistant or related genes. For example, sly-miR160a and novel_mir_762 target several Auxin response factors. Auxin is not only an important plant hormone affecting plant development, growth and abiotic stress, but also a new functional molecular to attenuate pathogens virulence (Robert-Seilaniantz *et al.* 2007; Spoel *et al.* 2008; Krishnamurthy *et al.* 2017). Although the high throughput sequencing technology used to characterize the sRNA component did not enable an accurate quantitative evaluation, our bioinformatics analysis combining northern blot confirmed several novel miRNAs conferring FOL infection in tomato, which offered us with an exciting further direction to investigate the role of miRNAs in resistance to FOL in tomato.

To conclude, by integrative analysis, our broad genome transcriptome RNA-seq/sRNA-seq data provide a comprehensive overview of the gene expression profiles between two different tomato cultivars Moneymaker and Motelle treated with FOL. Our results identified several disease resistance related genes/miRNAs which will facilitate further analysis of putative molecular mechanism of resistance in tomato upon to FOL, which eventually lead to improvement of Fusarium wilt disease resistance in tomato. It remains to be determined whether or how these candidate pathogen-related genes and miRNAs confirmed by qRT-PCR and northern blot respectively are overexpressed or knock-outed in Moneymaker/Motelle plant leading to the Fusarium wilt disease resistance. In this scenario, we would expect that overexpressing of these candidate pathogen-related genes or knock-outing of miRNAs would lead to enhance resistance to FOL and shed more light on exploring novel disease resistant genes/miRNAs in tomato.

## ACKNOWLEDGEMENT

This work was supported by grant of JSNSF: BK20161330, Jiangsu Province, China.

## ONLINE RAW DATA OF SEQUENCING

All raw data of sequencing were deposited at https://www.ncbi.nlm.nih.gov/bioproject/PRJNA407898

